# Comparison between ribosomal assembly and machine learning tools for microbial identification of organisms with different characteristics

**DOI:** 10.1101/2022.09.30.510284

**Authors:** Stephanie Chau, Carlos Rojas, Jorjeta G. Jetcheva, Mary Markart, Sudha Vijayakumar, Sophia Yuan, Vincent Stowbunenko, Amanda N. Shelton, William B. Andreopoulos

**Affiliations:** Department of Computer Engineering, San Jośe State University, San Jośe, CA, USA; Department of Computer Science, San Jośe State University, San Jośe, CA, USA; Department of Plant Biology, Carnegie Institution for Science, Stanford, CA, USA

## Abstract

Genome assembly tools are used to reconstruct genomic sequences from raw sequencing data, which are then used for identifying the organisms present in a metagenomic sample. More recently, machine learning approaches have been applied to a variety of bioinformatics problems, and in this paper, we explore their use for organism identification. We start out by evaluating several commonly used metagenomic assembly tools, including PhyloFlash, MEGAHIT, MetaSPAdes, Kraken2, Mothur, UniCycler, and PathRacer, and compare them against state-of-the art deep learning-based machine learning classification approaches represented by DNABERT and DeLUCS, in the context of two synthetic mock community datasets. Our analysis focuses on determining whether ensembling metagenome assembly tools with machine learning tools has the potential to improve identification performance relative to using the tools individually. We find that this is indeed the case, and analyze the level of effectiveness of potential tool ensembling for organisms with different characteristics (based on factors such as repetitiveness, genome size, and GC content).

**Author Summary:** Metagenomic studies focus on the challenging problem of identifying the presence and abundance of different species in a sample. This process typically involves the creation of digital reads from the sample which correspond to small parts of the genome sequence, and then have to be assembled together by a genome assembly tool. More recently, machine learning approaches have been applied to a variety of bioinformatics problems, and in this paper, we explore their use for organism identification, and how they might complement traditional bioinformatics approaches. We conduct experiments with two representative state-of-the-art machine learning approaches and six metagenomic assembly tools in the context of two synthetic datasets. We find that for organisms with certain characteristics (levels of repetitiveness, GC content, and genome size), ensembling metagenome assembly tools with machine learning tools has the potential to improve species identification performance relative to using the tools individually.

## 1 Introduction

Metagenomics studies can reveal the presence and abundance of different species in a sample, and can also be used to detect changes in organism composition across samples. For example, analyzing sequencing reads of the human microbiome can be used to uncover differences between individuals that may be indicative of the presence of a disease, and analysis of soil samples can capture the bacterial diversity at various time points or locations. Identification of species in metagenomics typically involves detecting the ribosomal 16S rRNA sequence and other marker genes. The ribosomal 16S gene is conserved and is used as a “fingerprint” that can identify the precise species or strain in a metagenomic sample. While numerous rRNA gene microbial surveys have shed light on the composition and characteristics of many ecosystems inhabited by bacteria and archaea [1–3], identifying precisely what organisms are in a sample continues to be a challenging problem. While existing metagenomic analysis methods have shown a reasonable performance at the genus and higher taxonomic levels, identifying closely related species and strains presents a significant challenge, because of the diversity of databases, genetic relatedness, and strengths or weaknesses of different tools [4]. In this paper, we focus on the problem of species-level organism identification from metagenomic samples, and explore the performance of both whole-genome sequencing tools, and state-of-the-art deep learning-based machine learning tools, with a focus on identifying the types of organisms (based on repetitiveness, genome size, and GC content), for which different tools perform better than others, as well as the potential improvement in performance that could result from ensembling these two types of tools.

We focus on whole-genome sequencing based on reconstructing the 16S rRNA gene from whole genome assemblies. We experiment with short reads, which are more affordable to produce than long reads, and have lower error rates. While long-read sequencing methods offer tools for correcting the errors with a consensus sequence analysis, the extra sequencing needed can be costly [5]. Long read sequencing requires high-quality DNA, which often increases the cost for metagenomics studies. Illumina short reads produce fewer sequencing errors with lower quality DNA and at a lower cost and are one of the most commonly used data formats for metagenomic studies.

In our study, we evaluated several commonly used metagenomic assembly tools, including PhyloFlash, MEGAHIT, MetaSPAdes, Kraken2, Mothur, UniCycler, and PathRacer, and compare them against against state-of-the art deep learning-based machine learning classification approaches represented by DNABERT and DeLUCS, in the context of two synthetic mock community datasets: *i.*) the mock community MBARC-26 consisting of 26 genomes [6], and *ii.*) a synthetic Hot Springs Mat sample dataset with 31 genomes of similar strains that we generated using the Camisim simulator [7] based on samples from Mushroom Spring and Octopus Spring, Yellowstone National Park [1–3, 8–10]. These datasets allow us to evaluate the effectiveness of the different approaches on genomes with a range of different characteristics along the dimensions of repetitiveness, genome size, and GC content.

Our analysis focuses on trying to understand whether ensembling metagenome assembly tools with machine learning tools has the potential to improve identification performance relative to using these tools individually. We find that the best performing tools in each class are phyloFlash and DeLUCS, and that they exhibit significant complementarity. In addition, we analyze the organism characteristics for which ensembling the tools has the most potential, including in terms of level of repetitiveness, genome size, and GC content. Our results indicate that an ensemble of genome assembly tools with machine learning approaches could result in improved species reconstruction, identification, and differentiation between similar strains.

## 2 Related Work

In this section, we will overview prior research on organism identification that focuses on short reads. There are two broad approaches for identifying the species and strains in short-read metagenomic data: *i.*) amplicon PCR sequencing followed by a read-level clustering or statistical analysis, *ii.*) whole genome shotgun sequencing followed by assembly. In the former case, we can categorize an Operational Taxonomic Units (OTUs) when 97% similarity thresholds is met [11, 12]. In the latter, the whole genome is assembled (including 16S rRNA), which gives us the ability to identify organisms at the species or strain level. Both approaches are followed by phylogenetic profiling. Shotgun metagenomic sequencing can identify microorganisms at the species or even strain level by profiling the variants, while 16S amplicon sequencing is generally limited to identifying microbes at the genus level due to insufficient discriminatory power of the read pairs from the V3/V4 region or sequencing errors [13–15]. Assembly tools are used to convert raw sequencing data into meaningful sequences that can identify the organisms in a sample precisely. However, assembling the 16S rRNA gene, which has a coverage higher than the rest of the genome, can be problematic for assembly tools. With the Silva and Kraken2 databases containing a majority of eukaryotic organisms and with less than 1% of existing microbes having been identified, identifying all strains is often impossible [16, 17]. In addition, whole genome shotgun sequencing often miss the marker genes that can be used for organism identification due to low-coverage of the genome. The genomes retrieved are often incomplete, having sequenced only a few thousand bases from some organisms, thus marker genes can not be recovered. Since using genes outside of 16S rRNA for phylogenetic profiling has to rely on any gene that can be assembled, it is often inaccurate and 16S is a standard for profiling.

### 2.1 Taxonomic profiling

Taxonomic profiling of rRNA sequences has been a popular approach for inferring the composition of complex microbial ecosystems. Taxonomy-based tools have been proposed, many of which are machine-learning based, though they generally go down to the genus level. Quantitative Insights Into Microbial Ecology (QIIME) and mothur have been widely used tools for taxonomic analysis, with MAPseq and QIIME2 being two recent alternatives. Almeida et al. compared the default classifiers of MAPseq, mothur, QIIME, and QIIME2 using synthetic simulated datasets comprised of the most abundant genera found in the human gut, ocean, and soil environments [18, 19]. QIIME2 provided the best recall and F-scores at genus and family levels. However, MAPseq showed the highest precision, with miscall rates consistently *<* 2%. Using the SILVA database generally yielded a higher recall than Greengenes.

Prodan et al. compared six bioinformatics pipelines for 16s rRNA amplicon analysis, and explore recall vs. specificity trade-offs. Generally these pipelines make predictions at the order or phylum level [20]. ADA2 offered the best recall, at the expense of decreased specificity. USEARCH-UNOISE3 showed a good balance between recall and specificity. QIIME-UCLUST produced large number of spurious OTUs, as well as inflated diversity estimation measures.

Wang et al. introduce the RDP classifier, a näıve Bayesian classifier, which provides taxonomic assignments from domain to genus, with confidence estimates for each assignment. The majority of classifications (98%) were of high accuracy [21].

Related to our work, Fiannaca et al. explore the effectiveness of using Deep Learning models in taxonomic classification for both amplicon and shotgun short-reads based on simulated bacterial datasets [22]. The work explores the use of deep learning methods in the context of several bacterial datasets, and uses a number of factors to characterize the properties of the organisms and how that impacts the performance of traditional and machine-learning-based metagenomic assembly approaches, especially in the context of strains of the same organism. Deep Learning methods performed better than the RDP reference classifier for bacterial identification at taxonomic levels from phylum to genus with both architectures. For instance, at the genus level, deep learning reached 91.3% of accuracy, while RDP classifier obtained 83.8% with the same data.

Liang et al. [23] present DeepMicrobes, a deep learning-based computational framework for taxonomic classification that is less hindered by the lack of a taxonomic tree for newly discovered species, which is required by current metagenomics tools.

They trained DeepMicrobes on genomes reconstructed from gut microbiomes and discovered potential novel signatures in inflammatory bowel diseases.

Lonear-Turukalo [24] evaluates the possibilities to identify clusters in a human microbiome based on taxonomic profiles. The results show that careful selection of the algorithm parameters and ensemble design are needed to ensure a stable clustering.

### 2.2 Reference genome based approaches

There has been considerable work on reference-based assembly with an aim to species and strain resolution, which often improves upon denovo assembly methods. These methods are generally limited by whether the exact organism’s reference exists in the database. Anyansi et al. [4] present an overview of recent methods based on assembly reconstruction and methods operating with or without a reference database for resolving strains within a species in metagenomic samples. They confirm that identifying close strains in metagenomes is more challenging, due to the diversity of databases and genetic relatedness. They find that tools have different strengths and weaknesses relative to a sample that will affect performance, and propose a workflow for selecting which tool to employ considering the organism properties, such as genome complexity.

Zeng et al. [25] report reference-based ribosome assembly (RAMBL), a computational pipeline that combines a 16S rRNA reference gene database with metagenomic shotgun sequences. RAMBL reconstructs full-length 16S gene sequences from metagenomic sequencing data accurately and could identify low and high abundance microbes in mock, human and soil communities [25].

Cepeda et al. [26] present another method for reference-guided assembly of metagenomes, which reduces the computational cost of mapping reads to reference genomes through a sample-specific indexing, resolves ambiguous read mappings and improves human sample assemblies.

### 2.3 Machine learning approaches

Besides reference-guided assembly, there has been work on improving the assembly of metagenomes using machine learning, often toward an aim to improve species and strain resolution. An early work was iMetAMOS for whole-genome assembly, an automated ensemble assembly pipeline that selects a single assembly from multiple assemblies [27]. They demonstrated the utility of the ensemble assembly on 225 previously unassembled *Mycobacterium tuberculosis* genomes, as well as a *Rhodobacter sphaeroides* benchmark dataset, and for identifying potential contamination.

Another early work was PERGA, which resolves branches in the assembly graph that are due to tandem repeats using a machine learning-based greedy-like prediction strategy. PERGA detects different copies of the repeats to resolve the branches and make the contig extensions longer, thus improving genome assembly [28].

Padovanu de Souza et al. review a variety of machine-learning methods for pre-processing of genomic datasets and post-assembly error detection [29]. They discuss application of ML during assembly, such as for identifying read overlaps prior to the construction of contigs, deciding whether to branch in de-Bruijn graphs, chimeric contigs, and to detect homologous chromosomes in diploid genomes. Greener et al. review general machine learning principles for biologists and techniques to improve genome assemblies, including deep neural networks for processing the assembly graph [30].

Quince et al. introduce STrain Resolution ON assembly Graphs (STRONG), which identifies strains de novo, from multiple metagenome samples [13]. STRONG performs coassembly and binning into metagenome assembled genomes (MAGs), and stores the coassembly graph. A Bayesian algorithm, BayesPaths, determines the number of strains present, their haplotypes and abundances. STRONG was validated using synthetic communities, showing they can generate haplotypes that match known genome sequences.

Harris et al. compared two different machine learning approaches to distinguish the taxonomic profiling among different metagenomic samples; one is a read-based taxonomy profiling of each sample and the other is a reduced representation assembly-based method [31]. Among various machine learning techniques tested, the random forest technique showed promising results as a suitable classifier for both approaches. Random forest models developed from read-based taxonomic profiling could achieve an accuracy of 91%.

Different sequence representation methods have also been proposed, which combined with the above approaches may improve species resolution. Woloszynek et al. explore using word and sentence embedding approaches for nucleotide sequences, which may be a suitable numerical representation for downstream machine learning applications (especially deep learning). The results show that embedding sequences results in meaningful representations that can be used for exploratory analyses or for downstream applications, such as binning [32–34]. Choi et al. present all-to-all comparison of metagenomes using k-mer content and Hadoop for precise clustering [35].

Ji et al. present DNABert, a model to generate genomic DNA embeddings inspired by the state-of-the-art Natural Language Processing (NLP) approach BERT [36, 37]. Natural languages have been shown to exhibit similar properties to DNA, such as polysemy (the coexistence of many possible meanings for a word or phrase). Similarly to the way BERT is used in NLP, DNABert is a pre-trained bidirectional encoder representation based on a transformer model, which can be further tuned to a specific task using an approach called transfer learning. This enables the application of deep learning to tasks for which lower amounts of data may be available. We evaluate DNABert as part of our experiments described in Section 4 as a representative of the state-of-the-art NLP approaches applied to genemic problems.

The work presented in [38] focuses on DeLUCS, an unsupervised approach for sequence classification based on deep learning. DeLUCS represents an important design point among the state-of-the-art machine learning approaches since is unsupervised and uses deep learning, and we evaluate it extensively in Section 4.

### 2.4 16S amplicon studies

Unlike whole-genome sequencing assembly, 16S amplicon studies focus on sequencing of marker genes, especially 16S rRNA. Often the goal of these studies is to assess differences between conditions, as well as linking back to whole-genome assemblies for profiling purposes. Amplicon studies assess the diversity of microbial species in a metagenomic sample by focusing on V3/V4 hypervariable regions of 16S rRNA genes using primers, instead of full length gene sequencing due to the limitations of short read sequencing platforms [39].

Amplicon studies group the reads into Operational Taxonomic Units (OTUs) to allow species identification, which first requires error correction of reads [40]. A common pipeline for 16S amplicon analysis starts by clustering sequences within a percent sequence similarity threshold (typically 97%) into OTUs. In each OTU one sequence is selected as a representative. This representative sequence is annotated and it is applied to all other sequences within that OTU. Such studies often find it challenging to determine the number of OTUs and their relative abundance in a sample, due to the difficulty of identifying population boundaries [41]. Since their focus is a portion of the 16S gene around the V3/V4 hypervariable region, they often have genus-level granularity and cannot identify the precise species or strains.

An alternative approach for amplicon analysis is the ASV approach, which is a statistical-based species identification method that goes the opposite way of OTU clustering methods. ASVs attempt to discriminate errors from true variants based on statistical methods, then mapping the resulting variants to databases of species to find the closest species. While with OTUs there is a risk of clustering different species into the same OTU if the threshold is too lenient, with ASVs there is a risk of artificially splitting a single genome into separate clusters [42–44].

Issues with amplicon studies involve the algorithms and parameters selected and the artifacts/errors of the sequencing. White et al. studied clustering methods for 16S rRNA sequences, showing that algorithm selection is critical to the quality of the results [45]. They provided a critical assessment of analysis methodologies that cluster sequences into OTUs showing that small changes in algorithm parameters lead to significantly varying results.

Nguyen et al. [46] focus on the problems of using sequence similarity for defining OTUs and discuss the shortcomings of this approach. They showed that the approach of using pairwise sequence alignments to compute sequence similarity results in poorly clustered OTUs. With OTUs annotated based upon a single representative sequence, poorly clustered OTUs can have significant impact on downstream analyses.

Mahe etl al. [47] determine that amplicon clustering methods have two fundamental flaws: arbitrary global clustering thresholds, and input-order dependency induced by centroid selection. They developed Swarm to address these issues by first clustering nearly identical amplicons iteratively using a local threshold, and then by using clusters’ internal structure and amplicon abundances to refine its results.

Schloss and Westcott [48] also investigate the parameters that affect the implementation of OTU-based methods. They show that it is impossible to create an accurate distance-based threshold for defining taxonomic levels and instead advocate for a consensus-based method of classifying OTUs. They propose a new heuristic that has a minimal effect on the robustness of OTUs.

Preheim et al. [41] determine two major challenges in calling OTUs: identifying bacterial population boundaries and differentiating true diversity from sequencing errors. They show that using the distribution of sequences across samples can help identify population boundaries even in erroneous data. They present a “distribution-based clustering” OTU-calling algorithm that uses the distribution of sequences across samples and demonstrate that its accuracy is competitive with other methods at grouping reads into OTUs in a mock community.

Huse et al. [49] examine deep sequencing of 16S amplicon libraries to facilitate the detection of low-abundance microbes in microbial communities. They found that a single-linkage pre-clustering methodology followed by an average-linkage clustering based on pairwise alignments more accurately predicts expected OTUs.

Edgar [50] tackles the high level of sequencing and amplification artifacts/errors found in amplified 16S sequences used to study microbial community structure. The UPARSE pipeline reports OTU sequences with *<* 1% incorrect bases in artificial microbial community tests, compared with *>* 3% incorrect bases commonly reported by other methods. The improved accuracy results in fewer OTUs, closer to the expected number of species in a community.

Hao et al. [51] proposed Bayesian clustering to tackle sequencing errors and profile the diversity of organisms, named “Clustering 16S rRNA gene sequences for OTU Prediction” (CROP). This method is useful to determine the number of OTUs and their relative abundance in a sample. CROP was robust against sequencing errors, which may provide a benefit for short and long-read sequencing methods.

Rasheed et al. [52] focused on the scalability of species diversity estimation by using locality sensitive hashing (LSH). The algorithm achieves efficiency by approximating the pairwise sequence comparison operations using hashing. They use LSH-based similarity function to cluster similar sequences and build groups of OTUs.

### 2.5 Statistical methods for organism identification

Besides OTU read clustering methods, there exist other statistical methods for identifying different microorganisms in 16S amplicon sequencing. Chudhary et al. [53] developed 16S Classifier using a machine learning method, Random Forest, for taxonomic classification of the short hypervariable regions of 16S rRNA sequence. On real metagenomic datasets, it showed up to 99.7% accuracy at the phylum level and up to 99.0% accuracy at the genus level. Tikhonov et al. [54] hypothesize that the standard approach to analyzing 16S sequence data, which relies on clustering reads by sequence similarity into OTUs, under-exploits the accuracy of modern sequencing technology.

They present a clustering-free approach to multi-sample Illumina datasets that can identify bacterial sub-populations, regardless of the similarity of their 16S sequences. Another approach puts sequences into bins based on their similarity to reference sequences (phylotyping), besides similarity to other sequences in the community (OTU clustering) [29, 48, 55].

There also exist statistical methods involving whole-genome shotgun sequencing of microbial communities. StrainFinder is a method that assigns strain genotypes and assesses strain frequencies for subpopulations of microbial communities with maximum likelihood estimates based on allele frequencies [56]. Ventolero et al. investigated 20 tools that attempt to infer bacterial strain genomes from shotgun read sequences [57]. They systematically evaluated six novel-strain-targeting tools on the same datasets and found that BHap, mixtureS and StrainFinder performed better than other tools. The performance of the best tools was still suboptimal though. Nayfach et al. [58] present the Metagenomic Intra-species Diversity Analysis System (MIDAS) pipeline for quantifying bacterial species abundance and strain-level genomic variation, including gene content and single-nucleotide polymorphisms (SNPs), from shotgun metagenomes. The method leverages a database of more than 30,000 bacterial reference genomes clustered into species groups. Subedi et al. propose a finite mixture of Dirichlet–multinomial regression models that accounts for the underlying group structure across gut microbiome samples and allows for a probabilistic investigation of the relationship between bacterial abundance within each inferred group [59, 60].

The results in the above work suggest that there is a need to move beyond simple clustering and assembly techniques for 16S identification and analysis. Our work expands on these by including additional tools in our analysis, both traditional bioinformatics tools and state of the art machine-learning-based tools. We included additional factors in our analysis of the genomes (e.g., coverage, repeats, GC) in order to get a deeper understanding of when different tools perform well and when they face challenges, especially at the species and strain levels.

## 3 Methodology

In this section, we describe the datasets used for our evaluation, the factors to characterize the data, and both the traditional bioinformatics tools and machine learning tools we used for organism identification. Our goal is to understand the differences in performance between the various tools when applied to organisms with diverse characteristics. We describe the datasets included in our study next.

### 3.1 Datasets

In order to explore and evaluate the performance of different metagenomic approaches for differentiating between species and strains, we utilized two datasets:

1. The mock community MBARC-26 benchmark dataset [6] contains 26 microbial organisms.
2. A simulated dataset we created to represent the widely studied Hot Springs Mat microbial environment from Mushroom Spring in Yellowstone National Park [1, 2, 8] contains 31 microbial organisms.

#### 3.1.1 Microbial Mock Community MBARC-26 Dataset

The mock community MBARC-26 [6], consists of 26 microbial organisms with finished reference genomes. The species contained in the dataset represent 10 phyla and 14 classes, and provide a range of GC content levels, genome sizes, and levels of repetitiveness. We chose to include this dataset in our study because it represents a diverse set of genomes, is a well-documented and widely used benchmark.

The organisms within the MBARC-26 dataset serve to provide a range of identification challenges. For example, this is visible in the Fig 1 and Fig 2 heatmaps, which shows the Blast identities for an all-to-all comparison of the 16S rRNAs of the organisms. There are organisms that are far apart in terms of BLAST ID percentage (*<*90%) and thus generally easier to differentiation from each other, there are also organisms of the same genus that are difficult to disambiguate (BLAST ID percentage *>*95%), and finally, there are organisms that are from different genera but have 90 *−* 95% similarity in terms of 16s composition, posing a significant identification challenge for standard metagenomic pipelines. We conduct the same analysis using a Hamming Distance percentage heatmap and observe a comparable distribution of 16s similarities across the organisms (Fig 1 and Fig 2).

**Fig 1.**
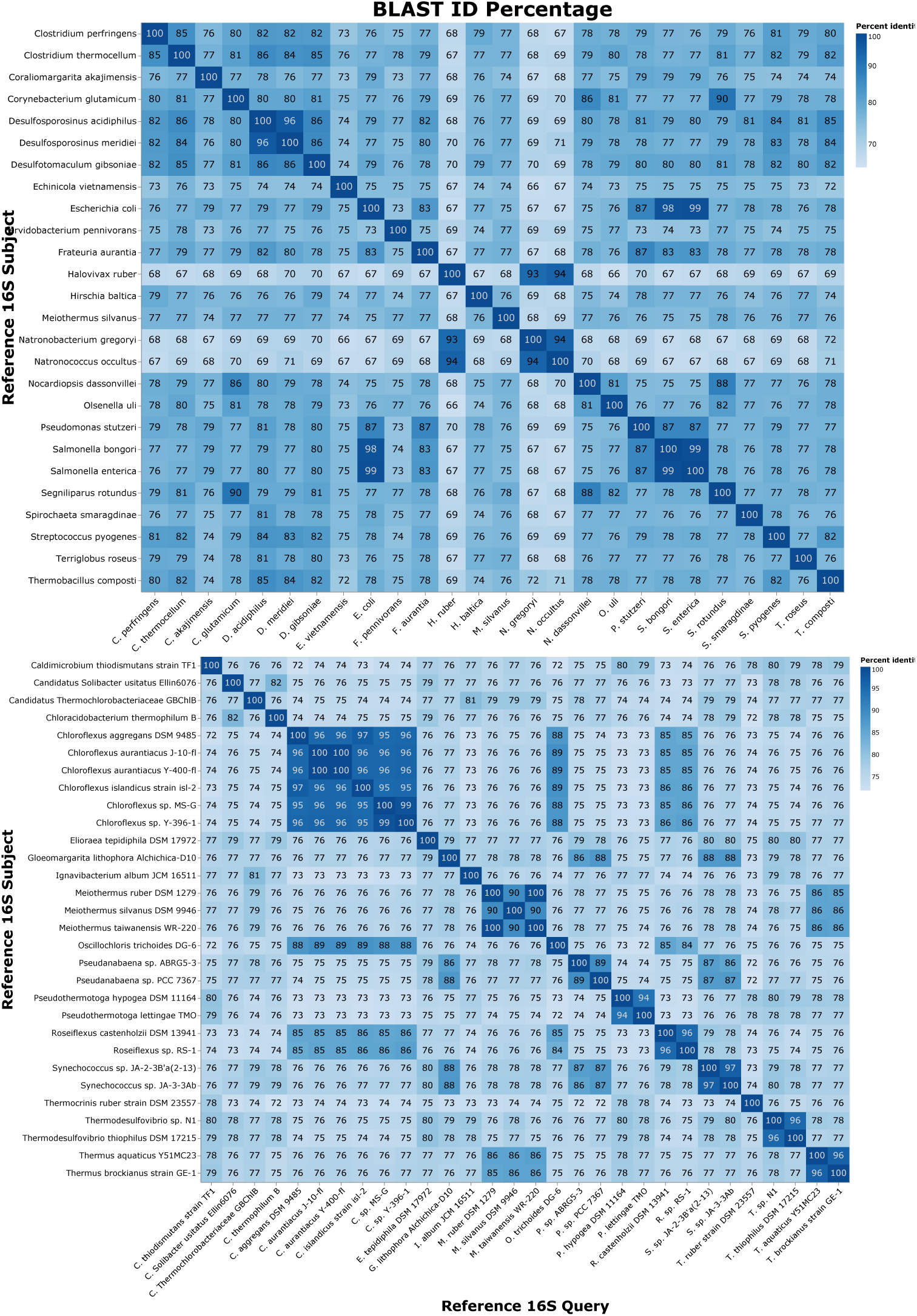
Blast identities between all vs. all of the 16S sequences in the MBARC-26 mock community dataset (top) and the synthetic Hot Springs Mat dataset (bottom). The matrices were generated with all-to-all Blast of the 16S sequences from the reference genomes. The similarity matrices between all of the species in the MBARC-26 mock community dataset show the diversity of the MBARC-26 dataset, but also clusters of similar species.

**Fig 2.**
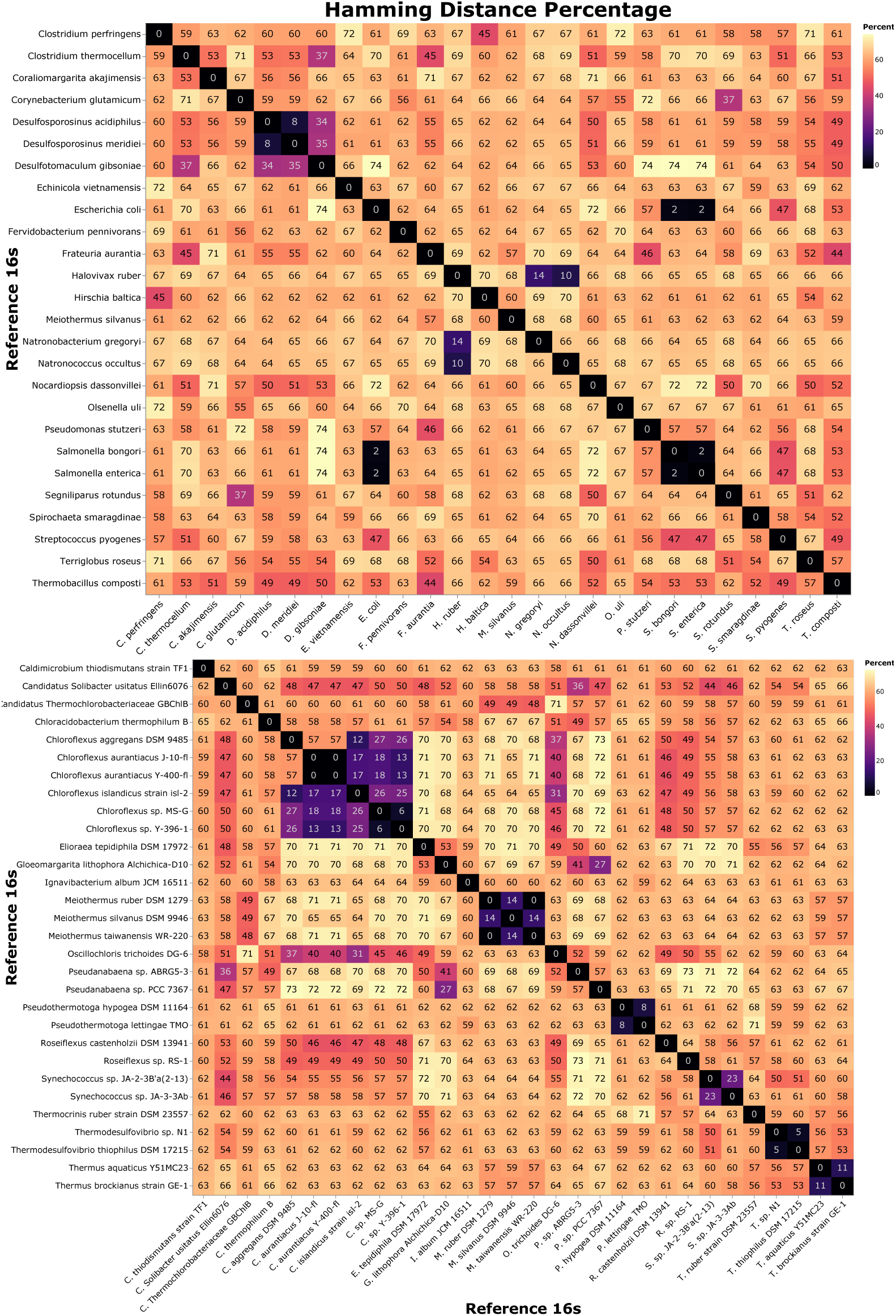
Hamming Distances between all vs. all of the 16S sequences in the MBARC-26 mock community dataset (top) and the synthetic Hot Springs Mat dataset (bottom). The matrices were generated with all-to-all Hamming Distances of the 16S sequences from the reference genomes. The similarity matrices between all of the species in the MBARC-26 mock community dataset show the diversity of the MBARC-26 dataset, but also clusters of similar species.

#### 3.1.2 Synthetic Hot Springs Mat Dataset

We created a synthetic Hot Springs Mat dataset by simulating reads from 31 microbial organisms with finished reference genomes detected in Octopus Spring and Mushroom Spring, Yellowstone National Park [1, 2, 8]. This is a widely studied environment with rich biodiversity and many unknown organisms, and is believed to hold the key to understanding important functional interactions between organisms over a range of environmental conditions and time frames. Reference genomes were chosen to represent closest available relatives to genera previously found in the mat and in an IMG metagenome assembly (SRA accession: SRP213403; IMG sample identifier: 3300032356). To simulate diversity, a number of reference genomes was chosen for some genera, including those sequenced from other hot springs (Supp. Table 1 – Hot Spring Mat Synthetic Metagenome Accessions).

**Table 1.** A comparison of both whole-genome and 16S assembly tools on the mock community MBARC-26. PhyloFlash achieved the most true positives (TPs) and few spurious hits that were not TPs. PhyloFlash has a built-in taxonomic assignment module.

|  | MEGAHIT<br>(Hits) | MEGAHIT<br>(TP) | MetaSPAdes<br>(Hits) | MetaSPAdes<br>(TP) | phyloFlash<br>(Hits) | phyloFlash<br>(TP) |
| --- | --- | --- | --- | --- | --- | --- |
| phyloFlash<br>Default config. | N/A | N/A | N/A | N/A | 23 | 22 (4 missing) |
| cmsearch with<br>Rfam 13.0 | 186 | 20 | 71 | 19 | 26 | 18 |
| Blast to IMG | 28 | 17 | 25 | 10 | 25 | 21 |
| Barrnap -outseq,<br>Blast with IMG | 28 | 17 | 27 | 15 | 25 | 21 |
| Kraken2 | 37 | 25 | 87 | 25 | - | - |
| DeLUCS | - | - | 26 | 11 | - | - |

|  | MEGAHIT<br>(Hits) | MetaSPAdes<br>(Hits) | phyloFlash<br>(Hits) |
| --- | --- | --- | --- |
| Barrnap -outseq | 17 Complete<br>& 21 Partial | 18 Complete<br>& 8 Partial | 45 Complete<br>& 1 Partial |
| Blast to SILVA | 1540 | 451 | N/A |
| Barrnap -outseq,<br>Blast with SILVA | 1540 | 192 | N/A |

The simulation data was generated with the CAMISIM simulator, which is widely used for generating synthetic genomic and metagenomic data [7]. We gave the NCBI IDs for each of the reference genomes to use as input to CAMISIM, which uses NCBI taxonomy for strain simulation and maps a reference genome to its NCBI entry using the NCBI ID. Relative abundance of the strains for the CAMISIM input (Supplemental Table 1) were made to reflect the relative abundances of genera reported in Liu et al 2011 [61] and an IMG metagenome assembly (SRA accession: SRP213403; IMG sample identifier: 3300032356). We simulated Illumina-generated short reads of length 150bp with a mean fragment size of 1000 and a standard deviation of 100 bases.

The resulting coverage is lower than for the mock community dataset as we wanted to limit the computation time of our experiments. The overall coverage amount does not impact our results – diversity of coverage between the organisms is what matters and what we have captured through this dataset. The resulting simulated metagenome contained 39.8M contigs with 6GB of data.

Based on the BLAST ID percentage similarity heatmap (Fig 1, Fig 2), we observe that the synthetic dataset contains more strains (*>*95%) than MBARC-26, and also more organisms that have 90% similarity and are thus challenging to disambiguate. These dataset characteristics are reflected in the Hamming Distance heatmap as well (Fig 1, Fig 2). Thus, we expect that the Hot Springs Mat dataset presents a more significant challenge to organism identification methods than MBARC-26.

### 3.2 Organism Characterisation

We used four factors to help characterize the organisms in our datasets: repetitiveness, genome size, GC content, and coverage. Each factor is computed for both the whole genome and the 16s sequence of each organism. We use the results to assess what types of organisms the various identification methods are able to differentiate well and which ones pose a challenge.

*Repetitiveness:* Bacterial genomes’ repetitiveness varies widely and is affected by the number of repeats and the largest repeats’ sizes [62]. Low complexity reads with high repetitiveness are considered particularly challenging as they tend to get mapped to the wrong species, e.g., during read alignment used in reference guided assembly approaches. Repetitive areas from a genome may interfere with the ability to assemble other genomes in the same metagenomic dataset. Thus, a genome’s assembly is affected by the dataset composition, as well as the individual genome’s repetitiveness. The assembly quality is affected by the collection of repeats from all organisms in a metagenome. For our datasets 1% *−* 3% repetitiveness is the medium range. We use the SPADE tool to measure repetitiveness of the genomes in our datasets [63].

*Genome size:* The genome size is computed by counting the number of bases in the reference genome. The average size of bacterial genomes is around 3 to 5 million base pairs, but there is a lot of variation around this average, with some genomes under 1M and others over 13M bases [64]. Genome size may pose a challenge for machine learning tools that employ long segments of the reference genome for species identification.

*GC content:* Segments of DNA with high or low GC content pose sequencing challenges to the lab instruments. Consequently, the data from DNA segments with high or low GC tends to have more errors in addition to being more repetitive. Assembling genomic areas of high or low GC content often contains mistakes. GC content varies widely among bacterial species, ranging from under 20% to over 70%. For most bacteria, the GC content falls within a relatively narrow range between 40% and 55%. E.coli’s GC content is 50.6% [64].

*Coverage:* Read coverage is a property to of the metagenomic sample, rather than the individual organisms in the sample. It plays a role in assembly as it may inhibit the assembly of certain species for which, the coverage is either too high (*>*150X) or too low (*<*30X). When the number of reads for a species is too low relative to other species in the sample, it may be difficult to assemble the sequence because of the shortage of reads needed to sufficiently map onto the reference genome. On the other hand, overly high coverage may cause de novo assemblers to throw out large groups of reads. Similar to repetitiveness, a genome’s assembly may be affected by the dataset’s collective coverage, besides the individual genome’s coverage. For our datasets 40*X −* 140*X* coverage is the medium range. We used the BBMap tool to measure coverage and genome size [65].

In our evaluation (Section 4), we denote the different ranges of each of the factors above as High, Medium, and Low, based on the thresholds described above.

### 3.3 Traditional Bioinformatics Tools

We evaluated the identification of metagenomic sequences in the context of our datasets with several state-of-the-art traditional bioinformatics assembly and identification tools, including phyloFlash [66], MetaSPAdes [67] and MEGAHIT [68]. We briefly overview each tool next.

PhyloFlash focuses on assembling the ribosomal 16S sequence, as opposed to the whole genome. It extracts all reads that match a ribosomal SSU database, assembles those reads using SPAdes [69] or MetaSPAdes [67] and then performs a taxonomic assignment (using the Silva reference database [70]). MetaSPAdes and MEGAHIT are whole-genome assembly algorithms that reconstruct longer segments of DNA (contigs) and involve a post-analysis to identify the ribosomes with cmsearch or barrnap and Blast. MetaSPAdes tends to provide more accurate results and takes into consideration the varying coverage levels within the datasets, and includes error correction, whereas MEGAHIT tends to be more efficient in terms of memory and time complexity. We used both tools to reconstruct the entire genome including the ribosomal sequence of each organism and experimented with several options for post-processing. The first option was the ribosomal database to select for Blast alignment. We experimented with both IMG [71] and Silva [70]. Another post-processing option we explored was to extract the ribosomal content with a profiler prior to Blasting. We experimented with both cmsearch and Barrnap (-outseq option), which are hidden-markov model (HMM) based profilers that detect sections of a genome sequence that match a ribosome followed by alignment to identify the correct species. We also evaluated Mothur, which compares sequences against ribosomal sequence databases Silva and RDP [70, 72] to identify the organisms [19]; as well as UniCycler [73] and PathRacer [74], which complement assembly tools to improve the result. UniCycler and PathRacer attempt to optimize the assembly graph using heuristics. We also evaluated Kraken2, which uses k-mers to match a query against databases of ribosomal sequences [16].

### 3.4 Machine Learning Tools

In this Section we describe the machine learning tools we used for organism identification. We explored the use of the unsupervised algorithm DeLUCS [38], which has been shown to work well on genomic classification tasks, and DNABert [36], an adaptation of state-of-the-art in Natural Language Processing deep learning and transfer learning approaches to the classification of genomic data. These algorithms are representative of a range of promising state-of-the art machine learning approaches and can help us gain insight into machine learning performance on different types of organisms, relative to traditional bioinformatics approaches. The machine learning tools employ the k-mers generated from input sequences.

#### 3.4.1 DeLUCS Algorithm

DeLUCS is an unsupervised approach and as such does not require the input sequences to be labeled for training the model, though it requires the number of clusters specified in training. DeLUCS starts out by generating a Chaos Game Representation (CGR) graphical representation of each DNA sequence of organisms it needs to learning to classify [75]. The CGR is a two-dimensional unit square image where the intensity of each pixel represents the frequency of a particular k-mer in the sequence. The k-mers from all organisms under consideration are used to generate each CGR image in order to ensure that images are comparable. The typical k-mer size used by DeLUCS is of length 6 (six-mers). Once the CGRs for a particular dataset of reference genomes is generated, the next key step in the algorithm is to generate mutant (mimic) sequences for each original sequence in order to imitate base mutations (transversions and transitions) that may occur in real life. Pairs of CGRs, one being the CGR of the original sequence and the other being one of the mimics, are grouped in batches and used to train a set of artificial neural networks (ANNs). The difference between the original versus mutant prediction in the output is used to represent loss, which the ANN tries to minimize. In particular, each ANN gets trained independently and has a softmax output layer, which outputs the probability that a sequence belongs to a particular class. The majority vote across the ANNs for each sequence is output as the final classification for the sequence. The taxonomic labeling and correspondence between clusters is a post-processing but essential component of DeLUCS. We used the known organism labels for our model evaluation. We used a threshold of 60% of a DeLUCS cluster mapping to a particular reference genome to identify a species as a hit. Additional parameter settings are enumerated in the Supplementary Information.

#### 3.4.2 DNABert Algorithm

DNABert is a deep learning-based pre-trained bidirectional encoder representation that can be used for downstream genomic tasks [36]. This approach was inspired by the BERT approach where a pre-trained language representation is created by training a deep learning model on a large general text corpus (e.g., news), and can then be trained further on smaller domain specific datasets, and applied to a variety of downstream tasks. This type of approach is referred to as transfer learning because a mathematical representation built based on one dataset/process is used effectively in the context of another dataset/process. This kind of transfer learning approach has been shown to perform well and is very common in deep learning and NLP due to the high compute requirements of training models on billions of documents from scratch. We use the DNABert pre-trained model and fine tune it by further training it with our data. We started out with the parameters recommended in [36], validated the key parameter choices with our data, and selected different parameter values than those recommended as needed.

#### 3.4.3 Data Pre-Processing

While the traditional bioinformatics tools were applied directly to raw reads, the amount of data contained in the raw reads dataset cannot be feasibly handled by a machine learning algorithm at this time. We conducted several experiments running the machine learning algorithms with samples of the raw reads (500-1000 reads per organism) but were not able to achieve reasonable performance and the tools were not able to handle a larger sample of reads.

DeLUCS has its own data pre-processing pipeline and generates k-mers directly from reference genomes. For DNABert, we generated 100– and 500– base pair shreds from the reference genome of each organism in our datasets, and used those to represent reads when using DNABert for classification [36]. We used 70% of the shreds for training and 30% for testing.

## 4 Results

In this section, we discuss the factor analysis on the organisms in our datasets and evaluate the performance of traditional metagenomic identification tools, comparing with the classification accuracy of newer machine learning tools.

### 4.1 Characterization of Genomes under Consideration

Both the MBARC-26 and the Hot Springs Mat datasets provide a diverse range of features as surfaced by the four factors we used in our study (repetitiveness, genome size, GC content, and coverage), thus providing a range of challenges for the identification and classification approaches we explore. Fig 5 to Fig 6 show the values for each factor for each dataset, and include whole genomes and 16S rRNA values.

One notable difference between the two datasets is the repetitiveness, which is up to 10% higher in the Hot Springs Mat dataset relative to the MBARC-26 dataset. We used a diverse set of genomes as shown in Fig 3 and Fig 4 heatmaps. The similarities and differences between genomes in the datasets provide different challenges for each of the tools we tested.

**Fig 3.**
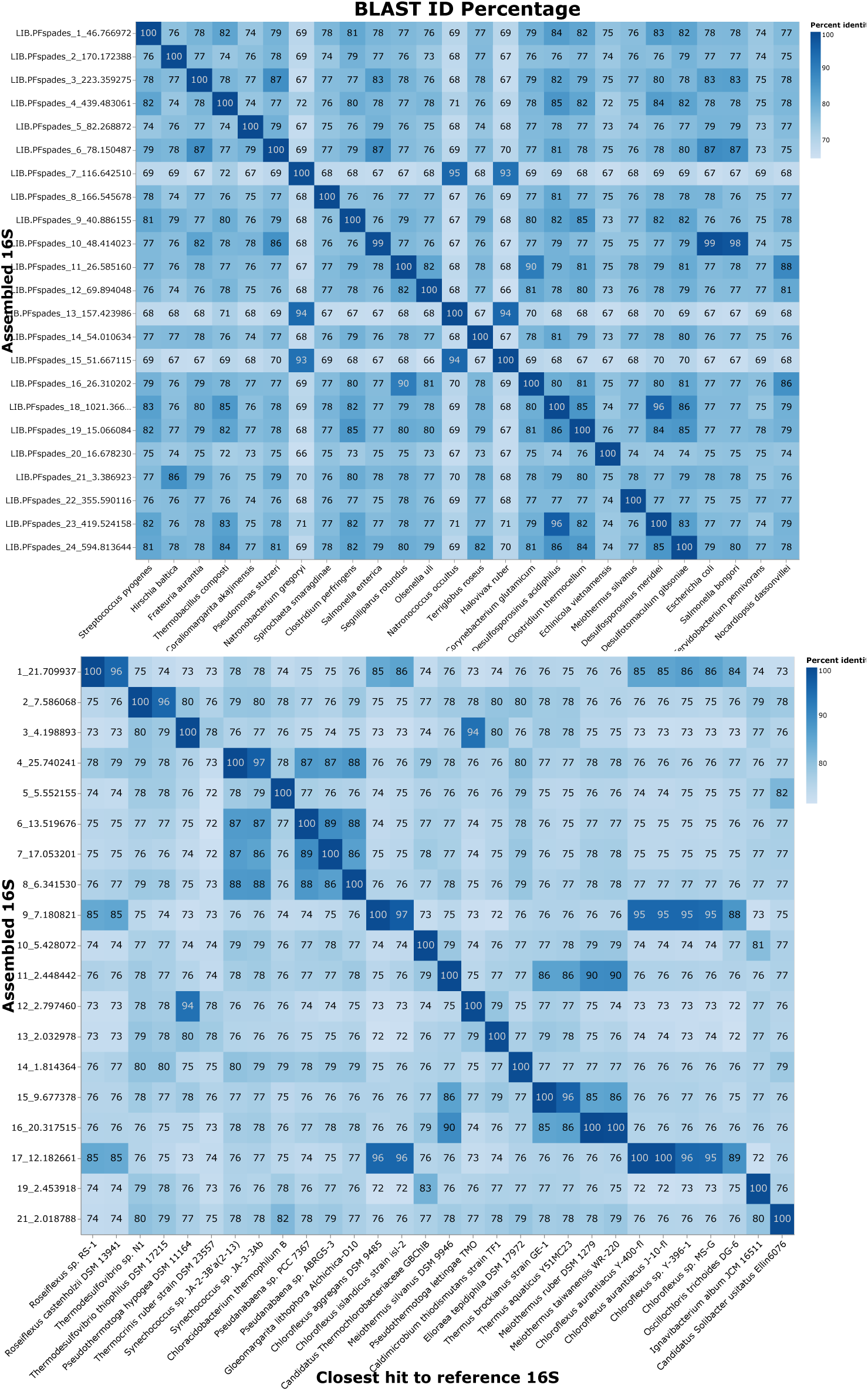
Blast identities of assembled vs. reference 16Ss for the MBARC-26 mock community (top) and the Hot Springs Mat Synthetic dataset (bottom). The matrices were generated with Blast of all assembled to all 16S reference sequences.

**Fig 4.**
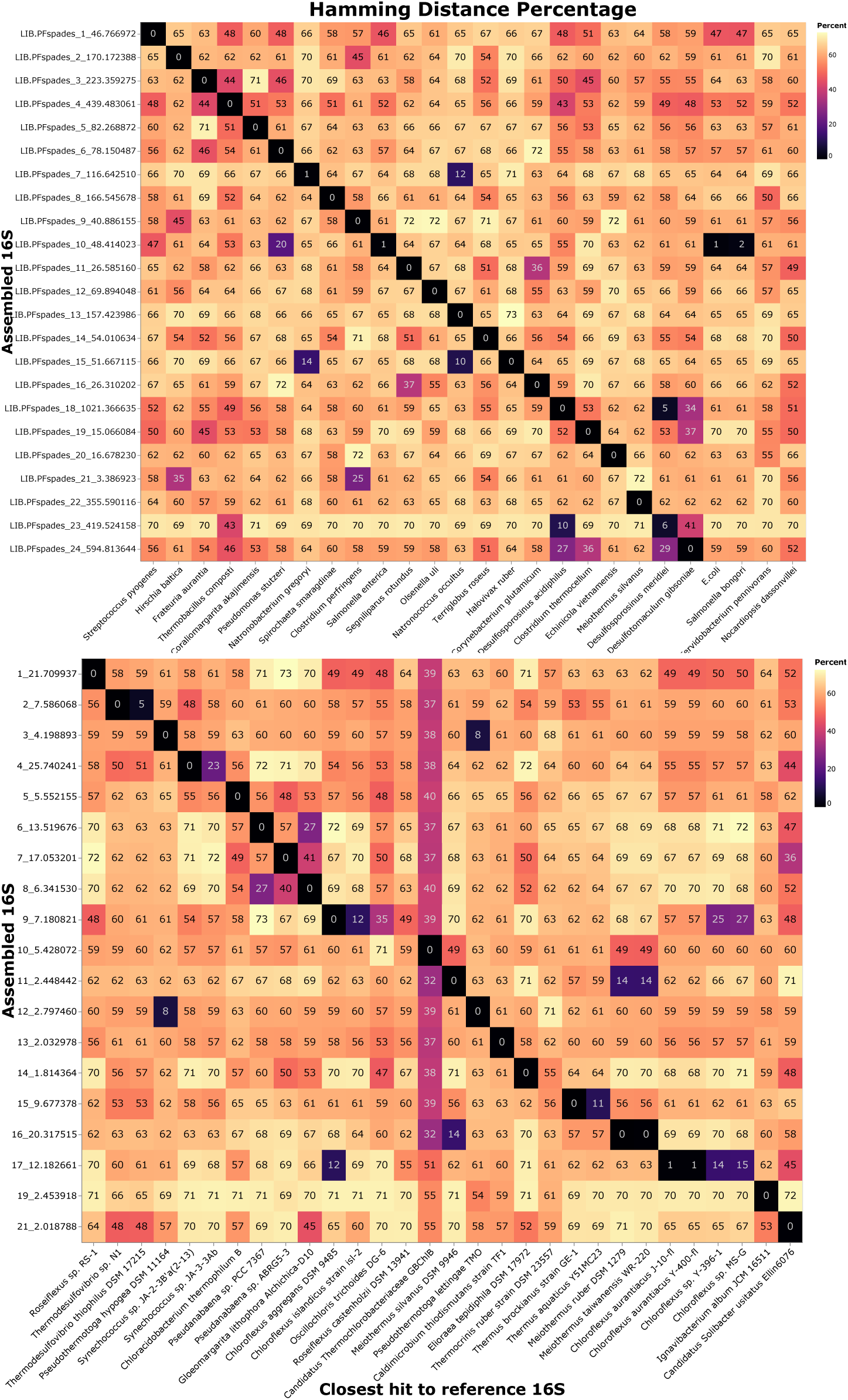
Hamming Distances of assembled vs. reference 16Ss for the MBARC-26 mock community (top) and the Hot Springs Mat Synthetic dataset (bottom). The matrices were generated with Hamming Distances of all assembled to all 16S reference sequences.

### 4.2 Metagenome Assembly Tool Performance

We focus our discussion on phyloFlash, MEGAHIT, MetaSPAdes, cmsearch, Barrnap, and Kraken2, since they performed the best amongst the traditional assembly tools that we evaluated. While we additionally tested PathRacer, Mothur, and Unicycler, they missed most of the organisms at the species level.

#### 4.2.1 Evaluation on MBARC-26 Mock Community

We evaluated the metagenomic assembly tools using two sequence alignment methods: cmsearch and BLAST, in conjunction with both the IMG and Silva rRNA databases.

Many of the metagenome assembly tools that we tested produced spurious hits, which are hits on organisms that are not within our dataset. Spurious hits are the total number of hits after subtracting the number of TPs. MetaSPAdes and MEGAHIT were used in conjunction with cmsearch, Barrnap, Kraken2, and BLAST (to IMG and Silva). Cmsearch outperformed Barrnap and BLAST; together with the metagenomic assembly tools, cmsearch gave fewer spurious hits and a slightly higher number of correct hits (true positives (TPs)). We summarized the results of applying phyloFlash, MEGAHIT, Kraken2, and MetaSPAdes on MBARC-26 in Table 1.

Our results show that the rRNA-focused assembler phyloFlash and Kraken2 perform better overall in terms of high TPs and few spurious hits. Kraken2 gave the most TPs with 25 organisms identified using the MEGAHIT or MetaSPAdes contigs, but it produced 12 spurious hits (for a total of 37 hits). Without any specific fine tuning relative to its default settings, phyloFlash was able to correctly identify 22 out of the 26 species with 1 spurious hit in the MBARC-26 mock community dataset. We tried to improve phyloFlash’s performance by augmenting it with the following three methods: Barrnap, cmsearch, or BLAST to IMG, but this did not lead to any improvement in the results.

PhyloFlash missed 4 species on the MBARC-26 mock community that did not get assembled: *Fervidobacterium pennivorans*, *Nocardiopsis dassonvillei*, *Escherichia coli*, and *Salmonella bongori*. There are several unique characteristics of these species, as shown in Fig 5-6, that might explain why they were missed by phyloFlash. *N. dassonvillei* is characterized by high GC content, high repeat content, and low whole genome coverage but high 16S coverage. *F. pennivorans* has high coverage and normalization of read coverage may result in a correct identification. *E. coli* and *S. bongori* both have low whole genome coverage but even though they have high 16S coverage, their similarity to *S. Enterica* (Fig 3 and Fig 4) makes their 16S reconstruction challenging.

Even though phyloFlash performed better than the whole-genome assembly tools, those tools combined with Kraken2 were able to successfully reconstruct the species that phyloFlash missed. This result can be seen in MEGAHIT and MetaSPAdes that identified the species *S. bongori*, *E. coli*, and *F. pennivorans*, which phyloFlash missed. Table 2 shows the missing species identified by cmsearch and Kraken2 for each output (MEGAHIT, metaSPAdes, and phyloFlash). This suggests that ensembling phyloFlash with some of the other tools may result in an improved overall reconstruction and identification. However, creating this kind of assembly may not be straightforward as cmsearch produced a large number of spurious hits in addition to having a high number of TPs (see Table 1)

**Table 2.** Missing species identified by cmsearch and Kraken2.

| Species | MEGAHIT | MetaSPAdes | phyloFlash |
| --- | --- | --- | --- |
| *S. bongori* | ✓ | - | - |
| *E. coli* | ✓ | ✓ | - |
| *F. pennivorans* | ✓ | ✓ | - |
| *N. dassonvillei* | - | - | - |

In the rest of this section, we perform a number of experiments to understand why phyloFlash was unable to identify the 4 species (*Fervidobacterium pennivorans, Nocardiopsis dassonvillei, Escherichia coli*, and *Salmonella bongori*).

#### Experiment 1: Are the missing species assembled?

PhyloFlash’s failure to reconstruct the four species in MBARC-26 could be due to one of two reasons: either it was not able to assemble the 16Ss, or the 16S was assembled but the species was not identified. To determine which of these is the case, we BLASTed the results of all the phyloFlash pipeline stages and we found that the four species were not detected at any stage. Next, we tested the mapping and assembly stages by running phyloFlash using reads from the four missing species, and determined that none of the four species were correctly assembled by phyloFlash. At the same time, although there were no assemblies, reads from the four species did map to phyloFlash’s reference database (using a read-based mapper called bbmap), showing that there were in fact reads from these species in the dataset. This helped us determine that the identification challenge for the species stemmed from the assembler (SPAdes) used by phyloFlash.

Next, we tried to analyze the reasons why the assembly failed. To do that, we investigated whether low coverage may explain the challenges faced by phyloFlash in assembling the organisms. When looking at the factor plots (Fig 5-6), we note that of the 26 species in the dataset, *N. dassonvillei* had the lowest coverage. The coverage for *E. coli* and *S. bongori* was also lower than average. On the other hand, all of their 16S sequences had high coverage as expected. In addition, one of the organisms, *F. pennivorans*, had the highest coverage of all 26 species. To understand whether the assembler issue was specific to the default phyloFlash assember (SPAdes), we conducted an experiment with an alternate assembler that we describe next.

#### Experiment 2: Exploring the use of a different assembler

Next, we ran phyloFlash on the whole dataset using the assembler MetaSPAdes instead of SPAdes, which is the default. We assumed that this would improve our results since MetaSPAdes is designed to assemble metagenomic data. While phyloFlash with SPAdes could not assemble *F. pennivorans*, phyloFlash with MetaSPAdes was able to recover this species in the final assembly. We believe that SPAdes underperformed because of the high coverage of *F. pennivorans*. However, phyloFlash with MetaSPAdes was still unable to assemble the other three species.

We ran phyloFlash with MetaSPAdes using the reads belonging to *E. coli, S. bongori*, and *N. dassonvillei*. Unlike when we used SPAdes, phyloFlash was now able to assemble *E. coli*. After removing the reads for *E. coli* and re-running phyloFlash with MetaSPAdes, *S. bongori* was assembled. However, when we removed the reads for both *E. coli* and *S. bongori*, phyloFlash was still unable to assemble *N. dassonvillei*. The low coverage of *N. dassonvillei* is a factor in the failure to assemble it. Only 3 out of the 399 reads for *N. dassonvillei* mapped to the reference database. This low coverage caused phyloFlash to abort before the assembly stage. Without sufficient alignment, the assembly phase did not have enough reads to reconstruct the whole sequence. Compared to SPAdes, MetaSPAdes may perform better when there is a wide variance in the coverage of organisms in metagenomes, but a low read coverage will cause issues.

#### Experiment 3: Analyzing repetitiveness

Our phyloFlash results indicate that low complexity (repetitive) reads tend to map to the same reference genome, even if the reads do not match the species. For example, phyloFlash could not assemble *S. bongori* and *E. coli* when both organisms are present. This suggested the presence of low complexity repetitive sequences. Therefore, if there are low complexity sequences in the data, some sequences may be mis-assembled. We used the repetitiveness tool SPADE [63] to check for unusually repetitive sequences and found that neither the whole genome nor the 16S sequences for any of the 4 missing species proved to be highly repetitive. Given these results, repetitiveness was unlikely to be the sole factor causing phyloFlash to fail for E. coli and S. bongori.

#### Experiment 4: Exploring the impact of organism similarity

Our final experiment focuses on organism similarity to understand the reasons for phyloFlash’s failed identifications. Assembling species that are very similar to one another is known to be especially challenging. Two species are very similar if they belong to the same genus and have a 16S rRNA sequence similarity of 97% or higher. High similarity between DNA sequences within a genus may challenge an assembler’s ability to accurately differentiate between distinct species or strains. Of the four species missed by phyloFlash, *S. bongori* and *N. gregoryi* are around 97% similar to *S. enterica* and *H. ruber* respectively. This level of similarity is at the threshold at which sequences may be considered as being from the same species. When MetaSPAdes replaced SPAdes, *N. gregoryi* was still missing. When combined with the reads for *Halovivax ruber* and *Natronococus occultus*, the 2 species most similar to *N. gregoryi* out of the 26 species, *N. gregoryi* was not assembled while the other 2 were. This confirms that organism similarity was one of the reasons phyloFlash failed in identifying some of the species.

#### 4.2.2 Evaluation on Hot Springs Mat Dataset

Results on the synthetic Hot Springs Mat dataset show that phyloFlash performed worse than on the MBARC-26 mock community dataset (Table 3). In particular, phyloFlash had 18 unique hits that matched one of the 31 organisms (TPs) and 17 spurious hits that did not match any of the 31 organisms. Kraken2 gave comparable TPs with 25 organisms identified using the MetaSPAdes contigs, but it produced 35 spurious hits (for a total of 60 hits).

**Table 3.** Results on the synthetic Hot Springs Mat dataset.

|  | BLAST to IMG |  | BLAST to Silva |  | Kraken2 |  |
| --- | --- | --- | --- | --- | --- | --- |
|  | TP | hits | TP | hits | TP | hits |
| MetaSPAdes | 22 | 592 | 0 | 4669 | 25 | 60 |
| MEGAHIT | 14 | 19 | 0 | 260 | 26 | 154 |
| phyloFlash | - | - | 18 | 35 | - | - |

MEGAHIT and MetaSPAdes produced a significant number of spurious hits on the Hot Springs Mat dataset, same as on the MBARC-26 mock community dataset. In particular, MetaSPAdes with Blast to IMG had 570 spurious hits and MEGAHIT had 5 spurious hits. When using Blast to Silva there were significantly more spurious hits.

The Hamming Distances and Blast identities indicated that the reconstructed 16Ss by phyloFlash were accurate at identifying the ground truth organisms – the Hamming Distances and Blast identities show a close similarity of the assembled 16Ss to the true organisms’ ribosomes (Fig 1, Fig 2) for both the MBARC-26 mock community and synthetic Hot Springs Mat.

Overall, we found that an rRNA-focused assembler like phyloFlash performed better than the metagenomic assembly tools followed by either Barrnap or cmsearch.

### 4.3 Machine Learning Performance

In this section, we present the results of applying DeLUCS [38] and DNABert [20] to our microorganism identification challenge in the context of the MBARC-26 mock community and our Hot Springs Mat datasets. We described the data pre-processing steps executed by DeLUCS and DNABert in Section 3. In the case of DeLUCS, we first ran its default pipeline taking as input the reference genomes. We also ran DeLUCS using MetaSPAdes contigs (500 per organism) and the 16S rRNAs extracted with Barrnap from contigs in order to evaluate its ability to classify from assembled ribosomal sequences. Both of the machine learning tools operate over the k-mers generated from the input genome sequences.

#### 4.3.1 DeLUCS Evaluation

DeLUCS converts the input sequences into k-mers, which are then used to generate Chaos Game Representation (CGR) representations [76]. In addition, one of the key parameters configured for DeLUCS is the number of mimics to generate during the training process – more mimics create more sequence variations to expose the algorithm to, but lead to higher computational overhead. To determine a reasonable setting for this parameter, we ran DeLUCS on the MBARC-26 dataset for 3-26 mimics (Fig 7) and found that using 5 mimics provides good performance (89% accuracy) at a reasonable computation load. The DeLUCS experiments in this section are based on using 5 mimics.

**Fig 5.**
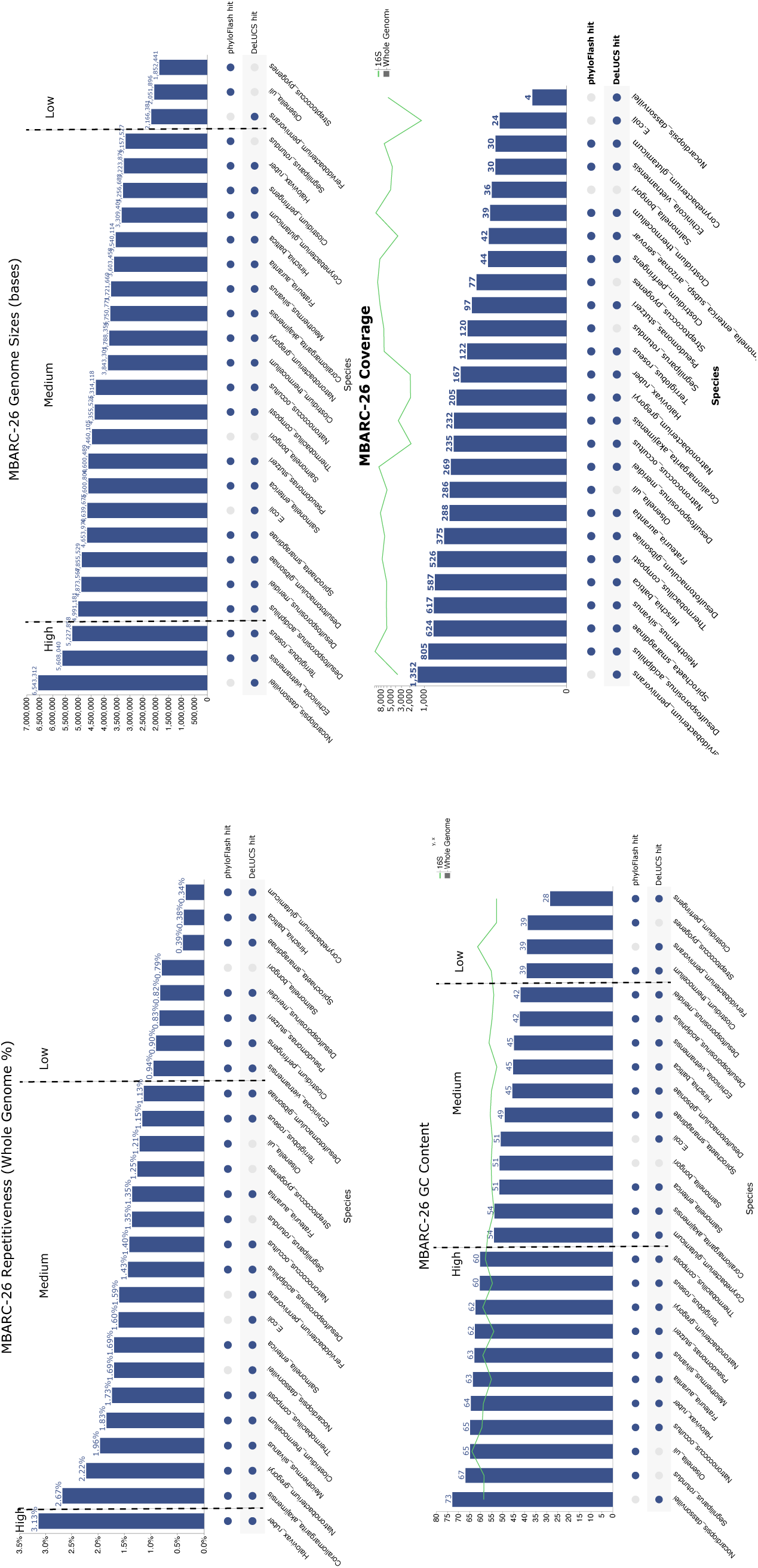
This figure shows the factor plots for the MBARC-26 dataset. The dotplot indicates which organisms phyloFlash vs. DeLUCS (on whole genome) missed or identified. The vertical dotted lines indicate which part of the data falls into the Low, Medium, or High range for each factor.

**Fig 6.**
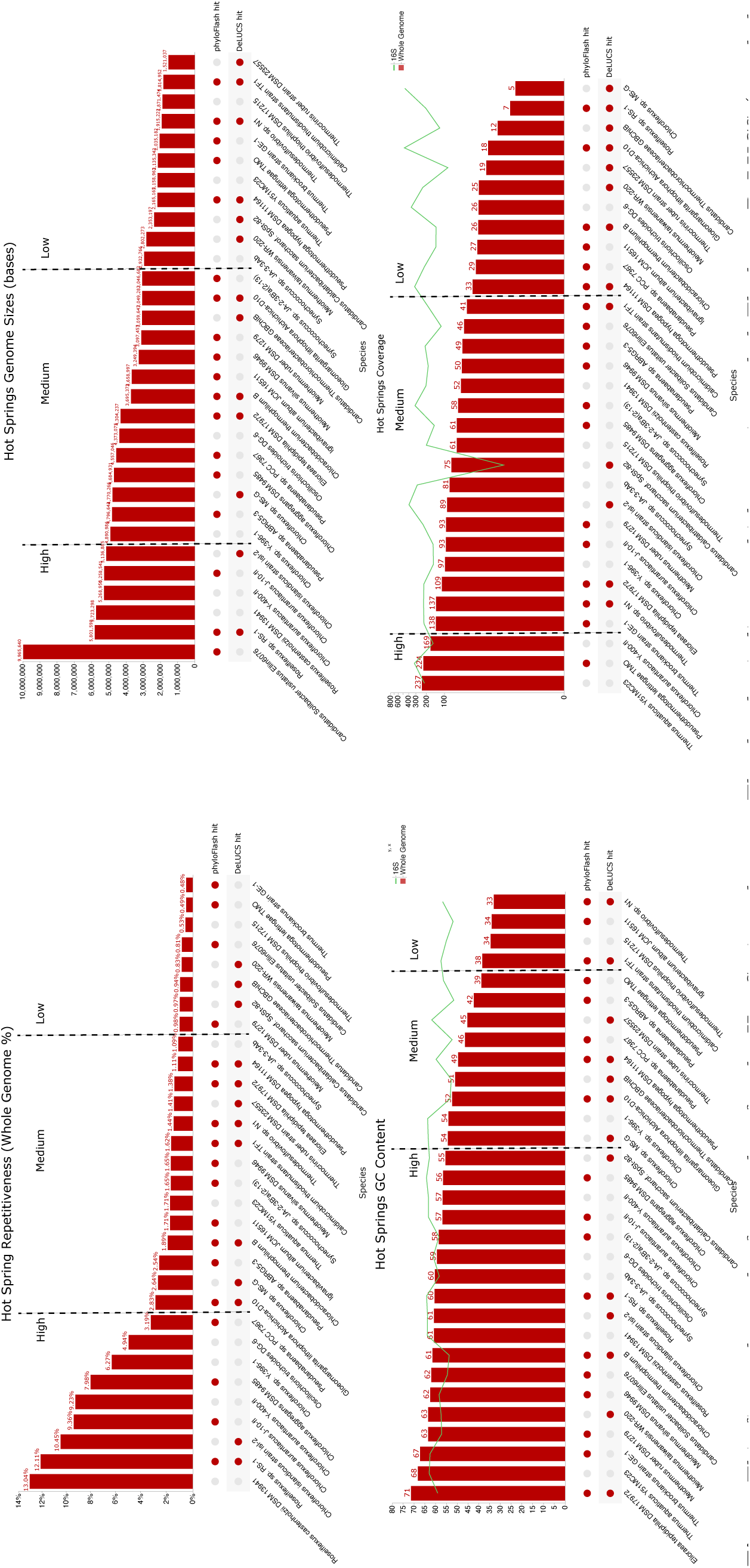
This figure shows the factor plots for the Hot Springs dataset. The dotplot indicates which organisms phyloFlash vs. DeLUCS (on whole genome) missed or identified. The vertical dotted lines indicate which part of the data falls into the Low, Medium, or High range for each factor.

**Fig 7.**
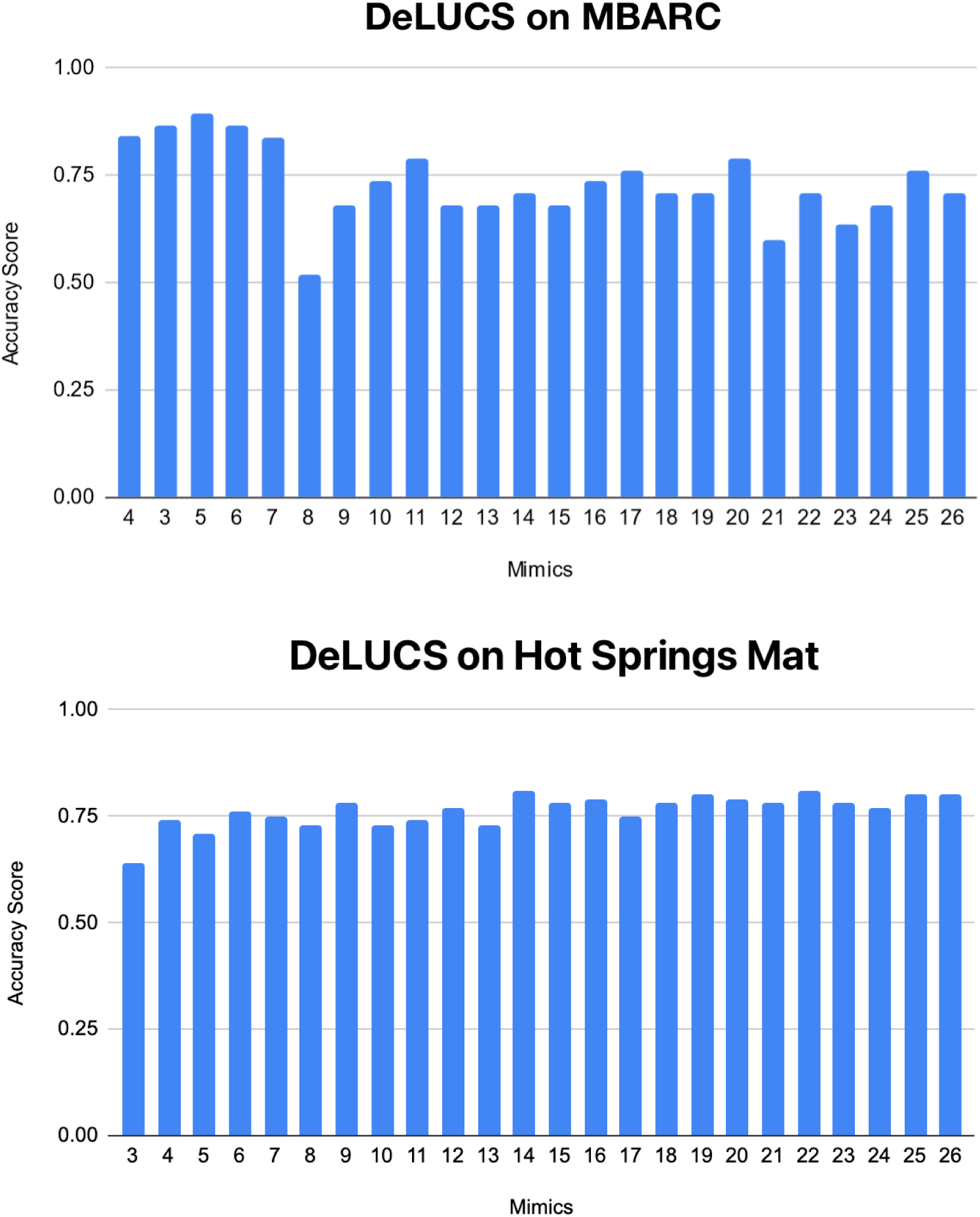
Accuracy achieved by different numbers of mimics in DeLUCS for MBARC-26 mock community (top) and synthetic Hot Springs Mat dataset (bottom).

DeLUCS achieved an accuracy of 89% on the MBARC-26 dataset, with 4 incorrectly predicted species: *O.uli, S.rotundus, S.pyogenes*, and *S.bongori*. *O.uli* and *S.rotundus* were falsely predicted to be *N. dassonvillei*, *S.bongori* was falsely predicted to be *E.coli*, and *S.pyogenes* was falsely predicted to be *D.gibsoniae*.

In terms of organism identification, we note that DeLUCS and phyloFlash have complementary performance in that DeLUCS is able to identify organisms that phyloFlash missed and vice versa. On the MBARC-26 mock community they each identified 22 out of 26 species and together they identified all but 1 organism (*S. bongori*).

Overall, DeLUCS had a lower performance on the Hot Springs Mat dataset than on the MBARC-26 dataset. This dataset was more challenging for phyloFlash as well.

The average accuracy when varying the number of mimics between 3 and 26 was 76% (Fig 7). The accuracy scores were mostly between 71 *−* 81% with the exception of the experiment where we used 3 mimics per sequence, which resulted in 64% accuracy. The upper half of the mimics (using 14-26 mimics) performed better than the lower half (all experiments resulted in *≥* 75% accuracy). Each additional mimic added around 2-4 minutes of time to the experiment on our infrastructure. We used 5 mimics for the results shown in the factor plots (Fig 5 to 6).

#### 4.3.2 DeLUCS Performance on rRNA from Assembled Contigs

While we were not able to evaluate DeLUCS directly on reads due to computational constraints (it could only handle 500 reads per organism which resulted in only 15% classification accuracy), we wanted to see if we might be able to use contigs as input instead. To that end, we ran DeLUCS on the Barrnap ribosomal output for the MetaSPAdes assembled contigs. For rRNAs extracted with Barrnap on the MBARC-26 mock community, the accuracy was about 44% for 10 mimics. For the Hot Springs Mat dataset, the accuracy was around 56% for 10 mimics. For MetaSPAdes contigs before ribosome extraction the classification accuracy was significantly lower at 25% for 3 mimics with 11 organisms out of 26 identified correctly (Table 1). DeLUCS’ use of mimics that are mutations of an original sequence shows the potential of machine learning tools to identify organisms that are mutations of known organisms and thus serve as a complement to tools like phyloFlash.

#### 4.3.3 DNABert Evaluation

We performed DNABert experiments utilizing GPU-powered nodes on our High Performance Cluster. Even so, we found that classification for both our datasets took 7-9 days to execute even for shreds of size 500, and resulted in poor performance (accuracy of 44% for MBARC-26 and 34% for the Hot Springs Mat data).

In order to determine if DNABert is able to handle at least binary classification experiments well, we conducted a number of experiments to determine what level of similarity of genomes it is able to handle. Fig 8 shows our results when using a structural similarity algorithm (SSIM) comparison between the CGR representations of the organisms’ 6-mer tokenized DNA sequences.

**Fig 8.**
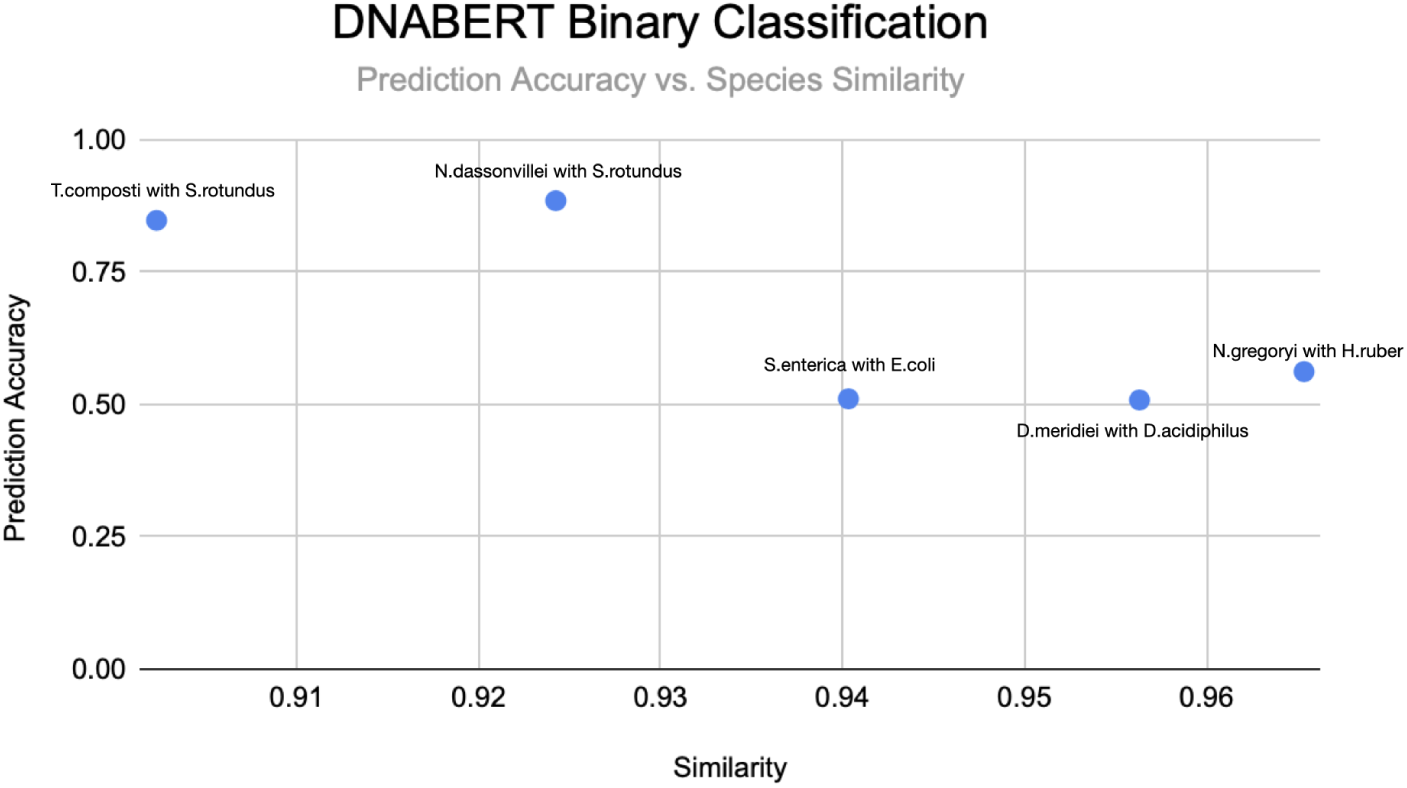
DNABERT classification accuracy for two species. As similarity between species increases the accuracy tends to decrease.

We found that while DNABERT is able to distinguish between two species when they are sufficiently dissimilar, it begins to produce poor predictions when species are at least 94% similar. For species pairs where similarity is 92.4% or less, DNABERT is much more accurate (88.5% compared to 51% prediction accuracy). Overall, we found that DNABERT is computationally cumbersome and underperformed with our data. As a result, we excluded it from the comparison with other tools in Table 1.

## 5 Discussion

To get deeper insight into the strengths and weaknesses of the best performing metagenomic identification tool in our experiments, phyloFlash, relative to the best performing machine learning tool in our experiments, DeLUCS, we look at their performance in the context of the factors we introduced in Section 3: repetitiveness, genome size, and GC content. We combined the two datasets into the same plot for each factor in order to facilitate our analysis (Fig 9 to 10). Because DeLUCS’ default pipeline generates k-mers from the input genome sequences uniformly (Section 3), we do not analyze the relative impact of coverage on the two tools.

**Fig 9.**
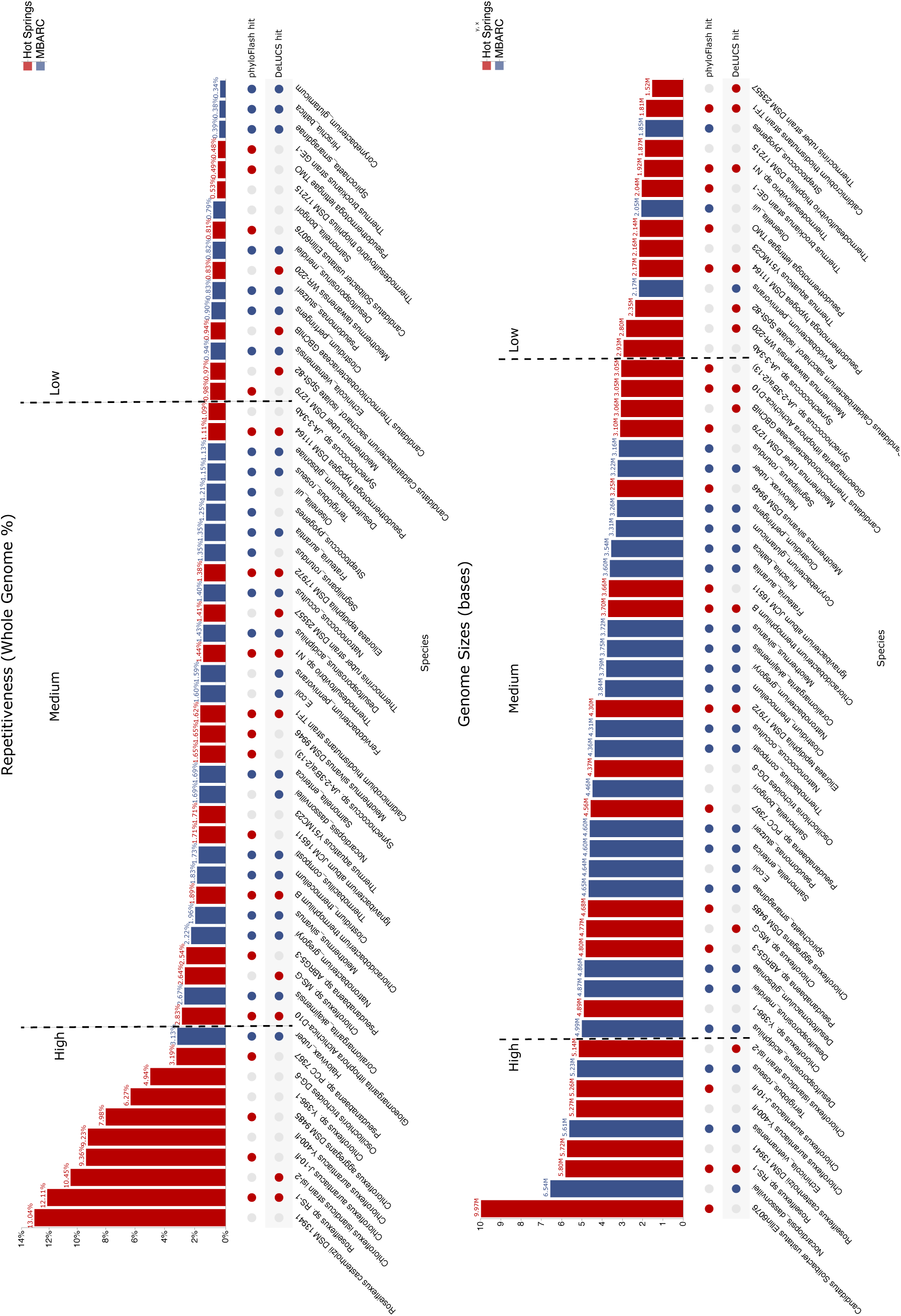
Merged factor plots of repetitiveness and genome size across the two datasets, showing which organisms were missed by one or both tools.

**Fig 10.**
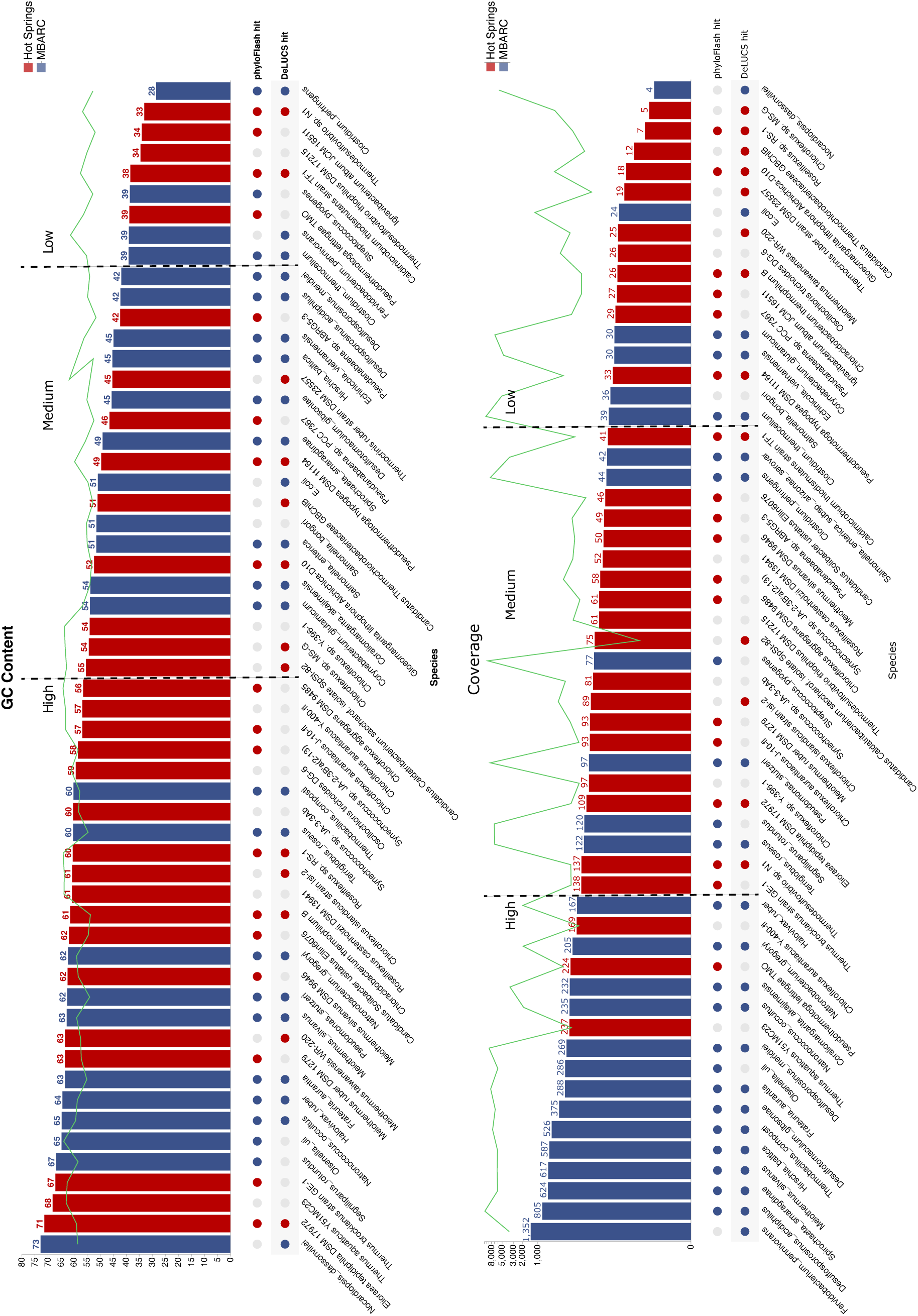
Merged factor plots of GC content and coverage across the two datasets, showing which organisms were missed by one or both tools.

In terms of repetitiveness, both phyloFlash and DeLUCS have challenges identifying organisms with high repetiveness, and to a lesser extent with low repetitiveness (Fig 9).

While phyloFlash performs better overall, the two tools have complementary performance – there are 23 species for which only one of the tools is correct. This indicates that 49 out of the 57 total organisms across the two datasets could have potentially been identified if the two tools were ensembled.

There are two instances where organisms with low repetitiveness (which should be easy to identify or classify) were missed by both tools. One such organism that presents a challenge to both tools is *Salmonella bongori*. The likely reason is that the tools are unable to differentiate it from the other Salmonella strain (*Salmonella enterica*) in the dataset. The other is, *T. theophilus*, which has a low genome size, which makes it challenging to identify and classify.

In terms of genome size (Fig 9), both phyloFlash and DeLUCS face challenges for organisms with high and low genome size. Understandably, shorter genome size organisms are harder to identify and classify. However, high genome size organisms should not pose a challenge unless there are other characteristics of these organisms that make their identification and classification challenging. We analyze further the two high genome size organisms that both tools are unable to recognize, *Roseiflexus* and *Chloroflexus a. Y-400*, and find that they are characterized by high-repetitiveness which explains why both tools failed.

Finally, we look at performance relative to GC content characteristics (Fig 10) and find that phyloFlash is most challenged by organisms with medium (but very close to the high threshold) and high GC content, whereas DeLUCS misses more organisms with high and low GC content. This indicates that the tools have complementary performance on organisms with medium and low GC content ranges.

To better understand when ensembling phyloFlash and DeLUCS would be most beneficial, we used the factor plots in Figures 9 and 10, to compute the ratio of organisms only one tool missed and organisms both tools missed relative to the Low, Medium and High range for repetitiveness, genome size and GC content (Table 4). These ratios indicate when the two tools are most complementary (bold ratio ratios in the table) and when they are least complementary (underlined ratios in the table) respectively.

**Table 4.** Complementarity between phyloFlash and DeLUCS is computed based on the merged factor plots. For each range (Low, Medium, High) of each factor (Repetitiveness, Genome Size, and GC Content), we show the ratio of organisms only one tool missed (complementarity):organisms both tools missed (non-complementarity). The ratios in bold indicate highest complementarity for a given factor, and the ratios that are underlined indicate the lowest complemetarity for a given factor.

|  | Low | Medium | High |
| --- | --- | --- | --- |
| Repetitiveness | 7:2 | **12:2** | <u>4:4</u> |
| Genome size | 8:3 | **11:3** | <u>4:2</u> |
| GC Content | **4:1** | 7:2 | <u>12:5</u> |

The highest complementarity between phyloFlash and DeLUCS is for organisms that fall within the medium range for repetitiveness and genome size, and the low range for GC content (highest ratio in each row of Table 4). Ensembling the two tools would be less effective for organisms with high repetitiveness, high GC content, or high genome size organisms (lowest ratio in each row of Table 4), with high repetitiveness presenting the biggest challenge. This indicates that additional tools need to be explored that can handle organisms with those characteristics.

## 6 Conclusion

In this paper, we explored state-of-the-art metagenomic assembly and machine learning tools in the context of metagenomic identification and found that these tools show significantly complementary performance. We started out with experiments with several representative tools in both classes, including the metagenomic tools, phyloFlash, MEGAHIT, and metaSPAdes, and the machine learning tools DeLUCS and DNABert, and conducted an in-depth comparison between the best performing tool in each class: phyloFlash and DeLUCS, respectively. We found that phyloFlash and DeLUCS exhibit significant potential for complementary performance. In addition, we analyzed the organism characteristics (repetitiveness, genome size, and GC content) for which ensembling the tools has the most potential. Our results indicate that an ensemble genome assembly tools with machine learning approaches could result in improved species reconstruction, identification, and differentiation between close strains. Finally, we note that state-of-the-art deep learning methods, such as those in DeLUCS and DNABert are currently largely opaque to interpretation, which makes it hard to get visibility into how and why the particular model arrives at its prediction. In the future, we plan to explore ways for enhancing or enabling the explainability of deep learning-based machine learning tools for organism identification.

## Acknowledgement

The research was funded by the College of Engineering of San Jose State University. Feqiao Brian Yu, Jacquelyn Stephanie Meisel, Rick Kim, Mihai Pop and Devaki Bhaya assisted in choice of relative abundances for the synthetic hot spring data set. The work (proposal: doi: 10.46936/10.25585/60001132) conducted by the U.S. Department of Energy Joint Genome Institute (https://ror.org/04xm1d337), a DOE Office of Science User Facility, is supported by the Office of Science of the U.S. Department of Energy operated under Contract No. DE-AC02-05CH11231.

## Supplementary Information

Data, figure images, and code notebooks are available under the Supplementary Information and under Zenodo. https://zenodo.org/record/7953871#.ZGp8fKXMKh8

